# CRISMER: A Transformer-based Interpretable Deep Learning Approach for Genome-wide CRISPR-Cas9 Off-Target Prediction and Optimization

**DOI:** 10.1101/2025.05.03.652008

**Authors:** Anamul Hoque Emtiaj, Rakibul Hasan Rafi, Muhammad Ali Nayeem, M. Sohel Rahman

## Abstract

CRISPR–Cas9 gene editing holds transformative promise for genetic therapies, but is hindered by off-target effects that undermine its precision and safety. To address this, we developed CRISMER, a hybrid deep-learning architecture that uses multi-branch convolutional neural networks to extract k-mer features and transformer blocks to capture long-range dependencies. This hybrid approach enhances the prediction and optimization of single-guide RNA (sgRNA) designs. CRISMER was trained on Change-seq and Site-seq datasets, using a 20 × 16 sparse one-hot encoding scheme, and evaluated on independent datasets including Circle-seq, Guide-seq, Surro-seq, and TTISS. CRISMER outperformed existing tools, achieving an F1 score of 0.728 and a PR-AUC of 0.818 on the CRISPR-DIPOFF dataset, and it generalized to fully independent datasets and to off-targets containing insertions and deletions. Ablation experiments confirmed that each architectural component contributes to performance, and CRISMER attained these results with substantially fewer parameters than transformer-and foundation-model baselines. It also demonstrated strong in silico specificity prediction and optimization capabilities, identifying sgRNA variants for targets such as PCSK9, BCL11A, and EXM1 with improved predicted off-target profiles. Interpretability analysis via integrated gradients confirmed the model’s focus on critical PAM-proximal regions and mismatch patterns. These results demonstrate that CRISMER significantly improves the accuracy and safety of CRISPR-Cas9, advancing its reliability for therapeutic applications.

## 1. Introduction

The CRISPR-Cas9 (Clustered Regularly Interspaced Short Palindromic Repeats associated protein 9) system has revolutionized genome editing by enabling precise genetic modifications across diverse organisms [12]. However, its clinical application faces a crucial challenge: off-target effects, where unintended genomic sites are modified due to sequence similarity with the target site [14]. These off-target modifications can lead to potentially harmful genetic alterations, making their accurate prediction and prevention essential for safe genome editing applications.

The development of computational approaches for CRISPR off-target prediction has progressed through three generations: empirical methods, traditional machine learning, and deep learning approaches [28]. Early empirical methods, including CFD [11] and uCRISPR [36], established foundational scoring systems based on experimental data. While these methods offered interpretable results, they struggled with complex sequence patterns and genome-wide predictions. CRISPROff [1] advanced this approach by incorporating energy models and biophysical properties, improving prediction accuracy while maintaining interpretability.

Machine learning-based approaches predict off-target activity using algorithms, such as random forest (RF) [3], support vector machine (SVM) [8], and XGBoost [6] by integrating various manually extracted sequence-derived features. CRISOT [5], for instance, utilized XGBoost in combination with molecular dynamics simulation-derived RNA-DNA interaction values to improve the predictive accuracy. However, the performance of machine learning-based tools varies significantly depending on the quality of constructed features, often necessitating substantial expert domain knowledge and manual curation to ensure relevance and effectiveness.

Deep learning approaches have significantly advanced off-target prediction through their ability to automatically learn complex sequence patterns. CNN-based architectures have been particularly successful, with several notable implementations. CnnCrispr [22] pioneered the application of convolutional neural networks for simultaneous local and global sequence feature extraction. DeepCRISPR [7] enhanced this approach by incorporating both sequence and epigenetic features through a CNN-autoencoder architecture. CNN_std [19] introduced position-specific nucleotide feature analysis.

Hybrid architectures have further improved the prediction accuracy. CRISPR-Net [20] combined CNNs with recurrent neural networks to capture both spatial and sequential patterns in sgRNA-DNA interactions. CRISPR-IP [37] integrated CNN, BiLSTM [29], and attention mechanisms [24] to focus on key sequence positions, while AttnToMismatch_CNN [21] specifically emphasized on mismatch positions between guide RNA and target DNA sequences.

Recent advances in transformer-based [34] architectures have pushed the boundaries of prediction accuracy. CrisprDNT [16] effectively handled variable-length input sequences, while CRISPR-BERT [23] adapted the BERT [9] framework to capture long-range dependencies in sgRNA-DNA interactions. CRISPR-DIPOFF [31] employed an RNN-based approach and leveraged the integrated gradients method [30] for interpretation purposes, thereby improving interpretability while maintaining high prediction accuracy. Most recently, the field has begun to explore foundation models: CCLMoff [13] repurposed a pre-trained RNA language model (RNA-FM) for off-target prediction, reflecting a broader trend toward transfer learning from large-scale biological corpora.

While existing tools have made significant progress, challenges remain in achieving both high prediction accuracy and effective sgRNA optimization. In this study, we introduce CRISMER, a deep learning–based pipeline for improved off-target prediction and streamlined sgRNA efficiency/specificity scoring and design. Table 2 summarizes how CRISMER differs from representative off-target prediction methods. Our key contributions include:

- We propose a hybrid architecture in which the two main components are matched to the structure of the prediction problem rather than simply stacked: a multi-branch convolutional block is designed to extract *k*-mer features at multiple scales, and a transformer block models long-range positional dependencies across the sgRNA-DNA duplex. A systematic ablation study (Section 3.2) confirms that both components and the multi-branch design itself, contribute measurably to performance.
- We introduce a 20×16 directional sparse pairwise encoding that preserves the orientation of each sgRNA-DNA base pair (e.g., distinguishing “AT” from “TA”), and we extend it to a 20 × 24 scheme that additionally represents single-base insertions and deletions. Ablation experiments show that this encoding outperforms conventional 4-and 5-channel one-hot schemes.
- We benchmark CRISMER against CNN-, RNN-, transformer-, and foundation-model-based methods under three complementary scenarios—a random split, fully independent datasets, and a mismatch-and-indel (bulge) setting—and report bootstrap-based uncertainty estimates together with a comparison of model size and training cost.
- We develop a comprehensive approach for sgRNA specificity evaluation and optimization, leveraging our deep learning-based model to generate guide RNAs with reduced off-target activity.
- We enhance model interpretability through Integrated Gradients, enabling transparent and biologically meaningful predictions focused on critical genomic regions.

## 2. Methods

### 2.1. Datasets

We assembled four groups of off-target datasets (Table 1), spanning a range of detection assays, guide designs, and levels of class imbalance. Groups I and II are taken from the aggregated benchmark released with CRISOT [5], in which every guide sequence is 23-nt with an NGG PAM. Group I (training) combines CHANGE-seq [18], an in vitro tagmentation-based assay that circularizes genomic DNA to measure genome-wide Cas9 cleavage, and SITE-seq [4], a cell-free biochemical assay that maps Cas9 cut sites in purified genomic DNA. Group II (independent evaluation) comprises CIRCLE-seq [32], a highly sensitive, reference-independent in vitro screen; GUIDE-seq [33], a cell-based assay that captures a short double-stranded oligonucleotide tag at Cas9-induced double-strand breaks; Surro-seq [25], a high-throughput assay that reads out indels across a pooled library of surrogate off-target sites; and TTISS [27], a parallelizable tag-integration assay for profiling many guides simultaneously. Group III originates from the DeepCRISPR collection [7] and is used to reproduce the CRISPR-DIPOFF [31] experiment; these guides retain their full 23-nt length with variable PAMs and permit up to six mismatches. Finally, Group IV is a mismatch-and-indel (bulge) benchmark curated by CRISPR-Net [20] from CIRCLE-seq-validated off-targets [32] and subsequently reused by CRISPR-BERT [23]; it contains off-targets with up to one single-base bulge in addition to mismatches, and we use it to assess generalization to indels.

**Table 1:**
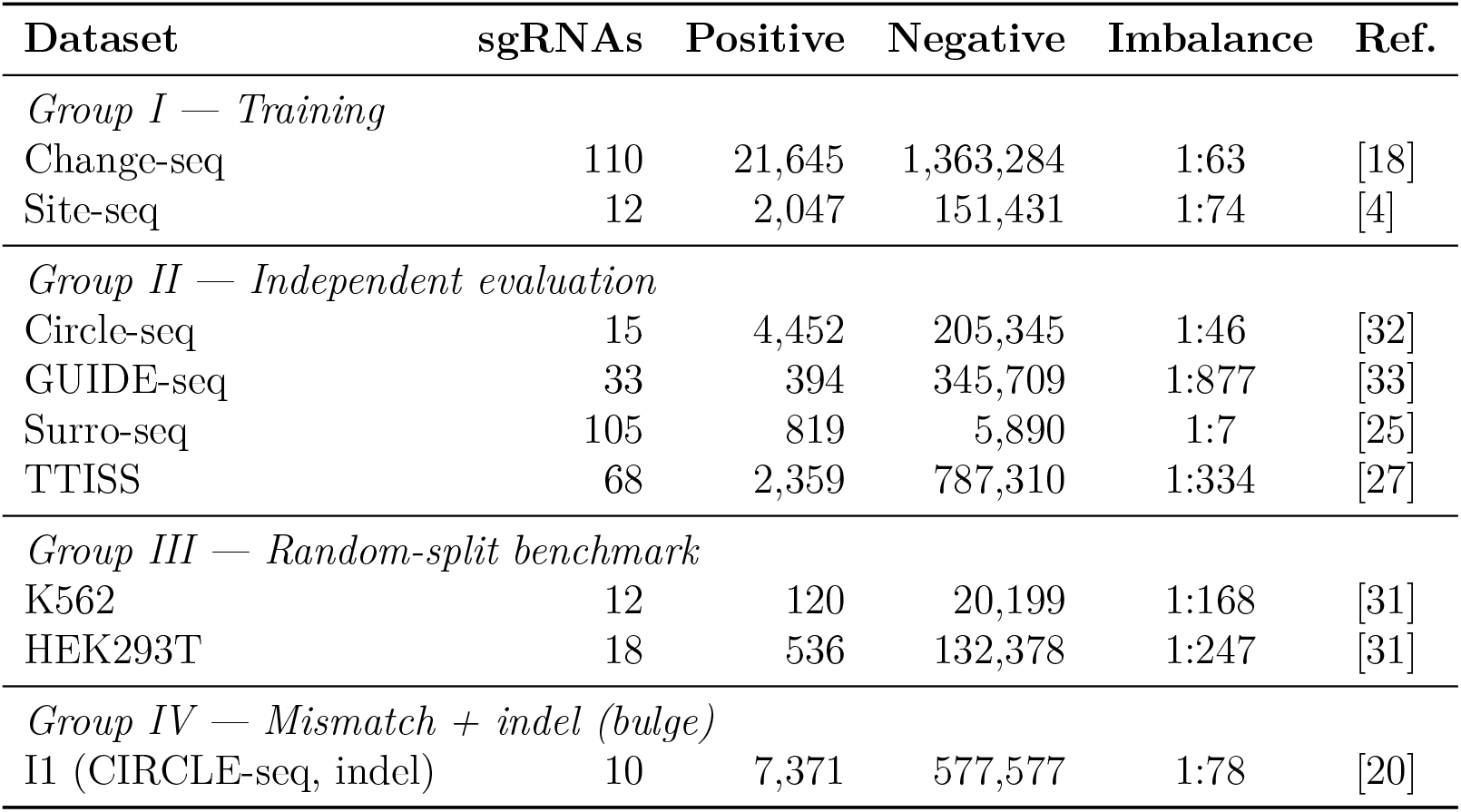
Overview of off-target datasets. Groups I (training) and II (independent evaluation) use 23-nt guides with NGG PAM from CRISOT [5]; Group III (random-split benchmark against CRISPR-DIPOFF [31]) uses 23-nt guides with variable PAMs; Group IV is a mismatch-and-indel (bulge) benchmark. The imbalance ratio is reported as the number of negative (inactive) sites per positive (active) site.

**Table 2:** Comparison of CRISMER with representative CRISPR-Cas9 off-target prediction methods. The “Indels” column indicates whether a method accommodates insertions/deletions (bulges) in addition to mismatches.

| Method | Feature extraction | Sequence model | Attention | Encoding | Indels |
| --- | --- | --- | --- | --- | --- |
| **CRISMER (ours)** | Multi-branch CNN | Transformer | CBAM + self-attention | $20 \times 16$ directional pairwise | Yes |
| CRISPR-Net [20] | Inception CNN | BiLSTM | — | Alignment-based pairwise | Yes |
| CRISPR-IP [37] | CNN | BiLSTM | Self-attention | 7-channel | Yes |
| AttnToMismatch_CNN [21] | CNN | — | Transformer | Learned embedding | No |
| CrisprDNT [16] | CNN | BiLSTM | Transformer | 14-channel | Yes |
| CRISPR-BERT [23] | CNN | RNN | BERT | Tokenized pair | Yes |
| CRISPR-DIPOFF [31] | — | BiLSTM | — | 4-/5-channel one-hot | No |
| CRISOT [5] | RNA-DNA fingerprint | XGBoost | — | 193 MD-derived features | No |
| CCLMoff [13] | RNA language model | Transformer | Self-attention | Pre-trained LM tokens | Yes |

All datasets are markedly imbalanced toward inactive (negative) sites, with negative-to-positive ratios ranging from roughly 7:1 (Surro-seq) to nearly 900:1 (GUIDE-seq). This pronounced imbalance motivates our choice of AUPRC as the primary evaluation metric.

### 2.2. Model

We developed a Convolutional Neural Network (CNN) and Transformer-based model to predict CRISPR-Cas9 off-target effects using a 20×16 sparse one-hot encoding scheme. Our proposed architecture consists of four main components: an input processing block, a multi-branch convolutional block, a transformer [34] block, and a dense network block (Figure 1). The model architecture integrates multiple attention mechanisms and parallel convolutional pathways to effectively capture both local and global sequence features while maintaining computational efficiency.

**Figure 1:**
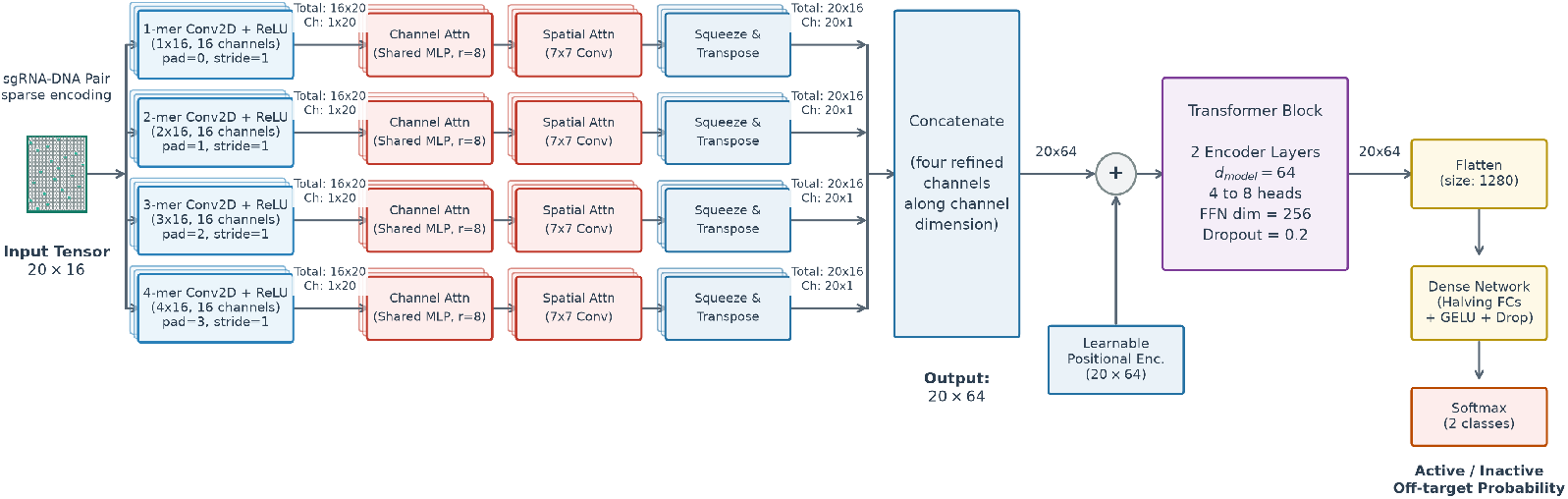
Detailed architecture of the CRISMER model for CRISPR-Cas9 off-target prediction. The model accepts a one-hot encoded input tensor of shape 20 × 16. Each parallel convolutional pathway is depicted as a 3D stack of 16 output feature channels. The annotations trace the tensor shape transformations from single-channel representations (shapes 1×20 and 20×1) to the combined branch tensors (shapes 16×20 and 20×16) before and after singleton dimension elimination and transposition. The branches are concatenated to yield a 20 × 64 feature map, which is combined with learnable positional encodings, processed by a Transformer block (comprising two encoder layers with *d*_model_ = 64), and mapped to final class probabilities via a fully connected feed-forward network.

#### 2.2.1. Input Processing Block

The input processing block converts each aligned sgRNA-DNA pair into a 20 × 16 sparse one-hot matrix. For each of the 20 spacer positions, the ordered pair formed by the sgRNA nucleotide and the corresponding DNA nucleotide is encoded as a one-hot vector over the 4 × 4 = 16 possible ordered base pairs, so that exactly one channel is active per position. Encoding the ordered pair rather than the two bases independently preserves directional information—for example, distinguishing an “AT” pairing from a “TA” pairing—which the conventional 4-or 5-channel one-hot schemes discard. For the mismatch-and-indel setting (Group IV), the pairing alphabet is augmented with eight gap symbols (A-, T-, C-, G-, -A, -T, -C, -G) to represent single-base insertions and deletions, giving a 20 × 24 encoding; the remainder of the architecture is unchanged except for the width of the first convolutional kernels. Furthermore, when the 3-nt Protospacer Adjacent Motif (PAM) is included in the sequence alignment (as in Groups I, II, and III), the sequence length increases to 23 nt. This expands the input dimensions to 23 × 16 for the mismatch-only configuration and 23 × 24 for the mismatch-and-indel configuration. In these configurations, the convolutional block preserves the sequence length of 23, and the learnable positional encoding is dimensioned to 23 × 64. Consequently, the output of the Transformer block has a shape of 23 × 64, which is vectorized into a 1472-dimensional representation before entering the fully connected layers. Because the representation is sparse and low-dimensional, it integrates efficiently with the downstream convolutional layers. Section 3.2 empirically compares this encoding against 4-and 5-channel one-hot baselines.

#### 2.2.2. Multi-Branch Convolutional Block

The multi-branch convolutional block employs four parallel pathways to capture k-mer features at varying scales, reflecting the biological relevance of k-mers in DNA sequences. Specifically, the first branch uses a 1×16 convolution with 16 output channels and no padding to extract 1-mer features, focusing on individual nucleotide patterns and their immediate context—crucial for identifying single-base preferences in CRISPR-Cas9 targeting. The second branch applies a 2×16 convolution with 16 output channels and a padding of 1 to capture 2-mer features, effectively analyzing dinucleotide combinations essential for local sequence composition and structural properties. The third branch utilizes a 3×16 convolution with 16 output channels and a padding of 2 to process 3-mer patterns, which often correspond to specific DNA structural motifs and binding preferences. Finally, the fourth branch implements a 4×16 convolution with 16 output channels and a padding of 3 to extract 4-mer features, capturing longer sequence patterns that may influence binding stability and specificity. Each branch takes a single input channel and produces 16 output channels. To ensure that every pathway preserves the input sequence length of 20 (or 23 when including the PAM), the convolutional kernel heights for the four branches are set to 1, 2, 3, and 4, respectively, with asymmetric zero-padding applied along the sequence dimension: the first branch uses no padding (total padding of 0), the second is padded at the end (total padding of 1), the third is padded at both the start and end (total padding of 2), and the fourth is padded with 1 row at the start and 2 rows at the end (total padding of 3). After a ReLU activation, each branch is refined by a Convolutional Block Attention Module (CBAM) [35]. To prepare the branch features (originally of shape 16 × 20 × 1 or 16 × 23 × 1) for concatenation, the singleton width dimension is eliminated to yield shape 16 × 20 or 16 × 23 (representing 16 feature channels). The tensors are then transposed to a sequence-first format of shape 20 × 16 or 23 × 16. Finally, the four branches are concatenated along the channel dimension to construct a 20 × 64 (or 23 × 64) feature map. The ablation study (Section 3.2) shows that the four-branch design outperforms three-and five-branch variants and that the attention modules provide a small but consistent gain.

#### 2.2.3. Transformer Block

The transformer block is employed to capture long-range dependencies in sequence data, a critical requirement for CRISPR-Cas9 off-target prediction. Unlike LSTMs or GRUs that process data sequentially, transformers leverage self-attention mechanisms to parallelize computations, significantly enhancing efficiency. This approach enables the model to simultaneously assess interactions across all nucleotide positions, which is essential given the global impact of off-target effects. Concretely, the block operates on the 20 × 64 feature sequence produced by the convolutional block, so the model dimension is *d*_model_ = 64. A learnable positional encoding—a trainable 20 × 64 parameter tensor—is added to this sequence to preserve positional order. The encoder is a stack of pre-normalization transformer encoder layers (layer normalization applied before each sub-layer), each comprising multi-head self-attention followed by a position-wise feed-forward network of dimension 256 (4 *×d*_model_) with a dropout of 0.2. Across the testing scenarios the encoder uses two layers and four to eight attention heads, selected per scenario; the complete search space and the per-scenario configurations are provided in Table S1 and Table S2 in the Supplementary Materials.

#### 2.2.4. Dense Network Block

The fully connected block maps the Transformer output to the final prediction. The 20 × 64 representation (or 23 × 64 when including the PAM sequence) is vectorized into a 1280-dimensional (or 1472-dimensional) vector, which is processed by one or two fully connected layers (with the depth optimized for each experimental scenario). Each hidden layer halves the feature dimension and applies a GELU activation followed by dropout (0.2) for regularization. A final linear layer produces two logits, and a softmax converts them into class probabilities for the active/inactive off-target prediction.

### 2.3. Training Configuration and Procedure

Model hyperparameters were selected by a genetic-algorithm search, with each candidate configuration evaluated by training the corresponding model and measuring its predictive performance. The complete search space and the optimal configuration selected for each testing scenario are provided in Table S1 and Table S2 in the Supplementary Materials. Using the selected configuration, we trained with the Adam optimizer [10] and applied a cosine-annealing schedule with warm restarts to improve training dynamics [15]. To counter the severe class imbalance, we used a weighted cross-entropy loss whose positive-class weight was also tuned by the search. Training was performed in mini-batches over randomly shuffled data.

### 2.4. Performance Metrics

We evaluated all models using the area under the receiver operating characteristic curve (AUROC), the F1 score, and the area under the precision-recall curve (AUPRC). While AUROC is widely reported, AUPRC is better suited to imbalanced datasets because it emphasizes performance on the minority (active) class; we therefore adopt AUPRC as the primary metric, with F1 as a complementary threshold-based measure. To quantify the uncertainty of each estimate and to enable meaningful comparison between models, we computed every metric over 1,000 bootstrap resamples of the corresponding test set and report the resulting mean *±* standard deviation; the corresponding 95% percentile bootstrap confidence intervals, alongside the mean *±* standard deviation, are provided in Tables S3–S6 in the Supplementary Materials. This bootstrap procedure yields confidence intervals for each metric, as recommended for benchmark comparisons on highly imbalanced data, and we consider a difference between two models to be reliable only when their bootstrap distributions are well separated.

### 2.5. Guide Efficiency Analysis

*Efficiency* refers to the probability of successful DNA cleavage at the intended genomic locus. Wild-type single-guide RNAs (sgRNAs) are known to exhibit high on-target activity; however, engineered sgRNA variants have been developed to improve specificity by reducing off-target effects, a trade-off that may in turn influence sgRNA efficiency. In this study, we propose a straightforward and reproducible approach for evaluating the efficiency of both wild-type and engineered sgRNAs at a defined target site.

To assess sgRNA efficiency, we employed our transformer-based off-target prediction model, which was trained on Group I data and independently evaluated on Group II datasets (Table 1) and which outperforms existing state-of-the-art methods. The output of the model’s final softmax layer, following temperature calibration, was used as a proxy for cleavage efficiency.

Deep learning models are often prone to overconfidence or miscalibration, particularly when handling complex or out-of-distribution inputs. To address this, we applied temperature scaling to the softmax output before its use as an efficiency measure. Specifically, a temperature parameter *T >* 1 was used to soften the raw logits, producing a smoother and more reliable score distribution. The resulting scores were subsequently min-max normalized to the [0, 1] interval to ensure comparability across the dataset.

To estimate the probability of active off-target effects, normalized confidence scores were binned, and for each bin we computed the proportion of off-targets classified as active relative to the total number of off-targets within that bin. This binned proportion served as an empirical estimate of the probability of an active off-target effect.

To assess the reliability of our model’s confidence scores relative to existing methods, we applied the same calibration procedure to several state-of-the-art deep learning-based predictive models; this comparison is presented in the Results section.

### 2.6. Guide Specificity Analysis

*Specificity* refers to the ability of a sgRNA to avoid inducing cleavage at off-target sites. A highly specific sgRNA minimizes activity at unintended genomic loci, thereby reducing the risk of undesired genetic modifications. To quantify specificity, we adopted the CRISOT-Spec scoring framework [5], which aggregates the predicted probabilities of all potential off-target sites into a single genome-wide specificity score. In our implementation, off-target probabilities are derived from our transformer-based prediction model rather than CRISOT’s original scoring function, and probabilities were weighted according to the results of our efficiency analysis such that sgRNAs with a higher likelihood of off-target cleavage are penalized more heavily. The aggregated off-target probability is defined as follows.

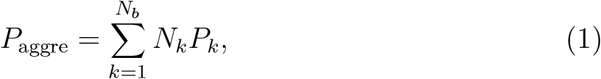

where:

- *N_b_* is the total number of confidence score bins,
- *N_k_* is the number of off-target sites in bin *k*, and
- *P_k_* is the estimated probability of cleavage for bin *k*, as derived from our model’s calibrated confidence scores.

The final specificity score is computed following the CRISOT-Spec formulation [5]:

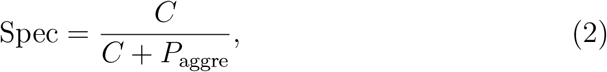

where *C* is a scaling constant. We retained the original formulation but re-calibrated *C* (set to 200 in our implementation) to keep specificity scores within an interpretable range given the score distribution produced by our model; all other aspects of the scoring scheme were left unchanged.

To evaluate genome-wide specificity, we analyzed all potential off-target sequences across the entire genome. Cas-OFFinder [2] was employed to identify candidate sites, allowing up to six mismatches between the sgRNA and the genomic sequence. Each predicted off-target was penalized based on its probability of being an active cleavage site, as defined in Equation (1), and the overall specificity score was then calculated using Equation (2).

### 2.7. Optimized Guide RNA Design

Although wild-type sgRNAs typically demonstrate sufficient on-target efficiency, they often exhibit significant off-target effects. For a given target site, our optimization process, CRISMER-Opti, begins with a wild-type sgRNA and introduces a limited number of mutations to create variants. Only sgRNAs with a confidence score above a predetermined threshold are further evaluated. Subsequently, we compute the genome-wide specificity score and select the sgRNA variant that maximizes this score.

When allowing a single mutation, each of the 20 positions in the sgRNA can be changed in three ways (excluding the original nucleotide), yielding 60 possible variants. For two or three simultaneous mutations, the number of variants grows combinatorially; however, additional mutations often reduce cleavage efficiency, so many variants fail to meet the confidence threshold. Consequently, our algorithm limits the number of simultaneous mutations and exhaustively evaluates all feasible variants to find the optimal balance between efficiency and specificity. Specifically, we select the variant with the highest specificity score among those that satisfy the confidence-score threshold.

### 2.8. Code, Environment, and Availability

All the deep learning models were designed using the PyTorch deep learning library [26]. The experiments were conducted in the Kaggle notebook environment, utilizing a single NVIDIA Tesla P100 GPU with 16 GB VRAM. We used the PyTorch Captum [17] library for model interpretation, which provides the implementation of integrated gradients. The datasets and source codes are available at https://github.com/Anamul-Hoque-Emtiaj/ CRISMER.

## 3. Results

### 3.1. Comparison of CRISMER with Previous Studies

In this section, we evaluate CRISMER against four representative off-target prediction models that span distinct methodological families: CRISPR-DIPOFF [31] (an RNN-based model), CRISOT [5] (a gradient-boosted model over physics-derived RNA-DNA interaction features), CRISPR-BERT [23] (a BERT/transformer-based model), and CCLMoff [13] (a recent foundation-model approach built on a pre-trained RNA language model). These models were chosen both for their established performance in prior studies and to cover the feature-engineering, recurrent, transformer, and foundation-model paradigms. To ensure a fair comparison, every competing method was retrained and evaluated by us under identical conditions—the same datasets, train/test splits, preprocessing, and 1,000-sample bootstrap procedure as CRISMER—rather than reusing numbers reported in the original publications. All metrics are reported as mean *±* standard deviation over the bootstrap resamples, and the ROC and precision-recall curves for representative trained models are shown in Figure 2.

**Figure 2:**
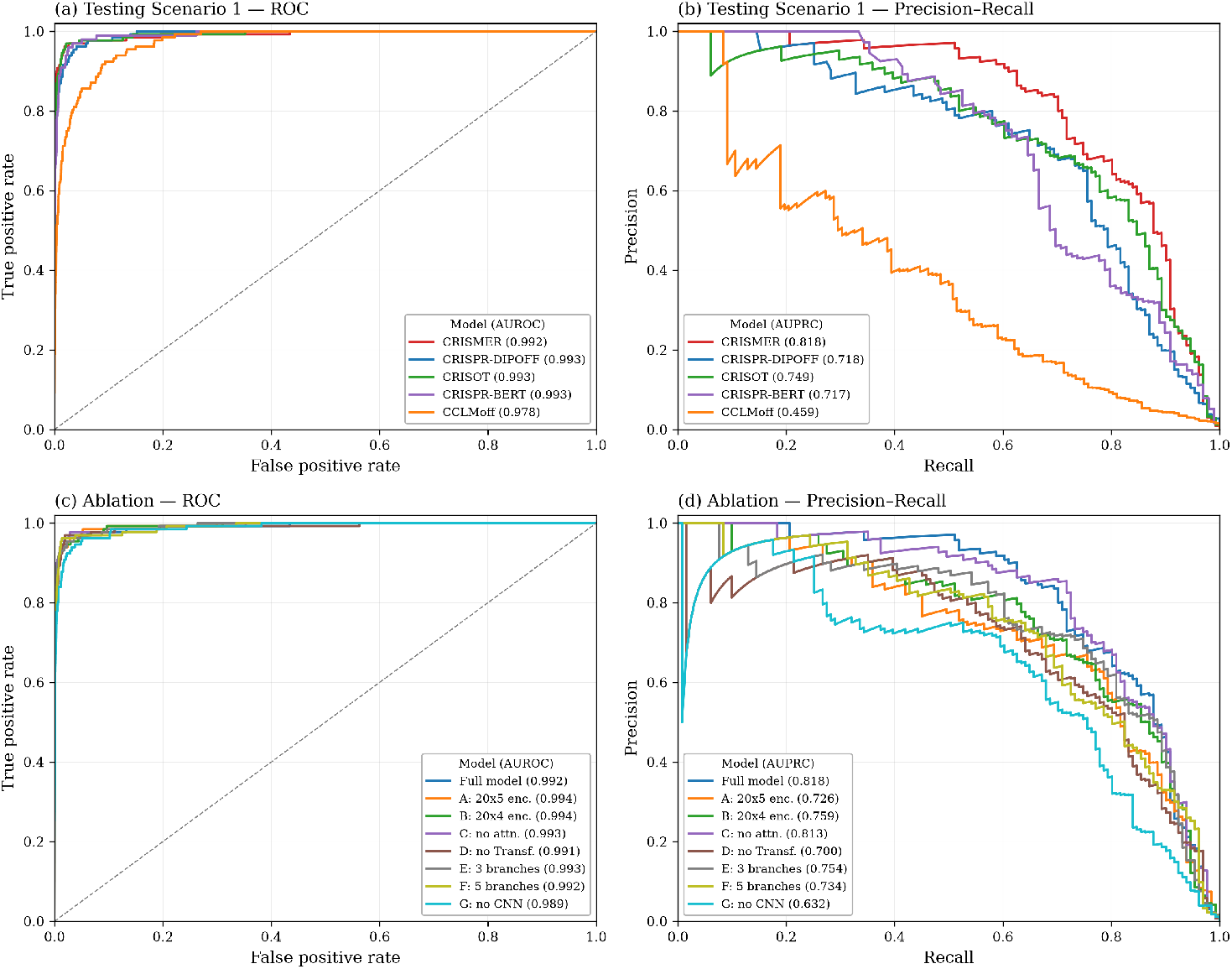
ROC and precision-recall (PR) curves for Testing Scenario 1 and the ablation study. (a) Testing Scenario 1, ROC; (b) Testing Scenario 1, PR; (c) ablation, ROC; (d) ablation, PR. Curves correspond to a representative trained model; the value in parentheses in each legend is the corresponding bootstrap-mean AUROC (ROC panels) or AUPRC (PR panels) reported in Tables 3 and 6.

**Table 3:**
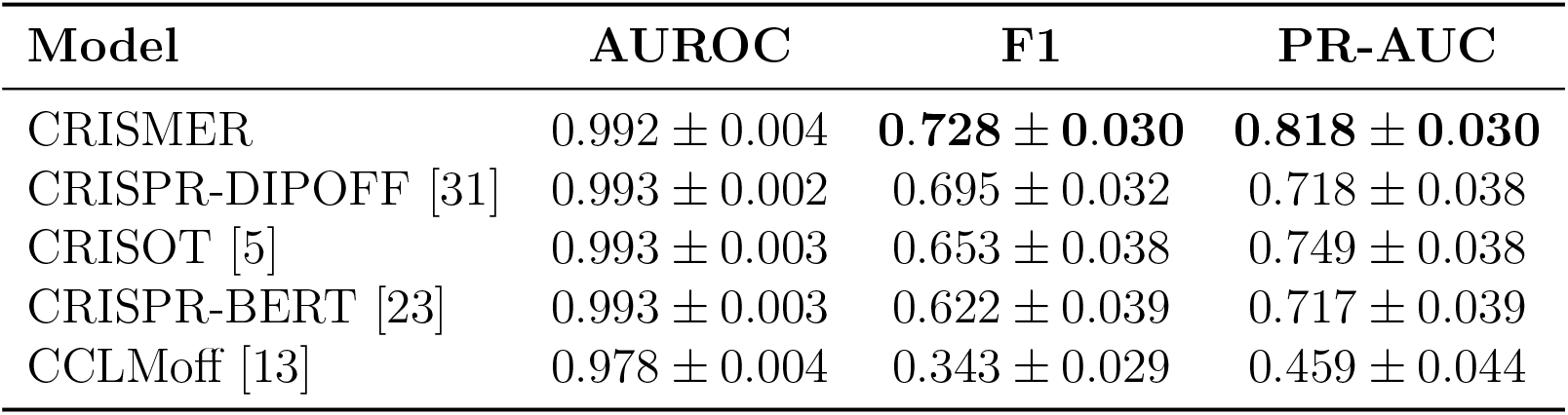
Performance comparison in Testing Scenario 1 (random 80%/20% split of the Group III data). Values are mean *±* standard deviation over 1,000 bootstrap resamples of the test set; the best mean F1 and PR-AUC are in bold.

| Model | AUROC | F1 | PR-AUC |
| --- | --- | --- | --- |
| CRISMER | $0.992 \pm 0.004$ | **$0.728 \pm 0.030$** | **$0.818 \pm 0.030$** |
| CRISPR-DIPOFF [31] | $0.993 \pm 0.002$ | $0.695 \pm 0.032$ | $0.718 \pm 0.038$ |
| CRISOT [5] | $0.993 \pm 0.003$ | $0.653 \pm 0.038$ | $0.749 \pm 0.038$ |
| CRISPR-BERT [23] | $0.993 \pm 0.003$ | $0.622 \pm 0.039$ | $0.717 \pm 0.039$ |
| CCLMoff [13] | $0.978 \pm 0.004$ | $0.343 \pm 0.029$ | $0.459 \pm 0.044$ |

#### 3.1.1. Testing Scenario 1: Random 20% as Test Set

The first testing scenario replicates the experimental setup of CRISPR-DIPOFF using the Group III data of Table 1, with an 80%/20% random train/test split. Table 3 reports the results. CRISMER attained the highest F1 score (0.728 *±* 0.030) and the highest PR-AUC (0.818 *±* 0.030)—the two metrics most informative under heavy class imbalance—ahead of CRISOT (F1 0.653, PR-AUC 0.749), CRISPR-DIPOFF (0.695, 0.718), and CRISPR-BERT (0.622, 0.717). The four sequence-based models achieved nearly identical AUROC (*≈* 0.99); AUROC is known to saturate on datasets with such extreme imbalance, which is precisely why we treat AUPRC as the primary metric. CCLMoff trailed substantially (F1 0.343, PR-AUC 0.459), a gap consistent with the large training-data requirements of pre-trained language-model approaches relative to the modest size of this dataset. The ROC and precision-recall curves in Figure 2(a,b) corroborate these rankings, with CRISMER dominating the high-precision region of the PR curve. Overall, CRISMER provides the best balance of precision and recall, consistent with the goal of reliable active-site detection.

#### 3.1.2. Testing Scenario 2: Independent Dataset Test (IDT)

In this scenario, the combined CHANGE-seq and SITE-seq data (Group I of Table 1) served as the training set, and the four Group II datasets were used only for evaluation. This is the most demanding setting, as the evaluation guides, assays, and genomic contexts all differ from those seen during training. Table 4 reports AUROC, F1, and PR-AUC. On the primary metric, PR-AUC, CRISMER achieved the best result on all four datasets: Circle-seq (0.654), GUIDE-seq (0.526), Surro-seq (0.466), and TTISS (0.436); notably, on TTISS it also exceeds CRISOT (0.403). The F1 ranking differs on the most extreme datasets: CRISOT obtained higher F1 on Surro-seq and TTISS, and CRISMER’s low F1 on GUIDE-seq (0.080) and TTISS (0.121) reflects the fixed 0.5 decision threshold applied under negative-to-positive ratios approaching 1:900, rather than poor ranking—its leading PR-AUC on the same datasets confirms that active sites are ranked highly. Moreover, TTISS is the largest of the Group II evaluation sets (Table 1), so the PR-AUC advantage there is the best supported by the data. CRISPR-DIPOFF underperformed across all datasets, while CCLMoff remained mid-ranked despite being trained on the largest available data. Overall, CRISMER ranks true off-targets more reliably than the competing methods across heterogeneous, fully independent datasets.

**Table 4:** Performance comparison in Testing Scenario 2 (independent-dataset evaluation; trained on Group I, tested on Group II). Values are mean *±* standard deviation over 1,000 bootstrap resamples. The best mean per metric within each dataset is in bold; PR-AUC is the primary metric.

| Dataset | Model | AUROC | F1 | PR-AUC |
| --- | --- | --- | --- | --- |
| Circle-seq | CRISMER | **0.973 ± 0.001** | 0.560 ± 0.006 | **0.654 ± 0.008** |
| | CRISPR-DIPOFF [31] | 0.869 ± 0.003 | 0.261 ± 0.008 | 0.346 ± 0.009 |
| | CRISOT [5] | 0.953 ± 0.002 | 0.253 ± 0.009 | 0.499 ± 0.009 |
| | CRISPR-BERT [23] | 0.972 ± 0.001 | **0.573 ± 0.007** | 0.620 ± 0.008 |
| | CCLMoff [13] | 0.949 ± 0.002 | 0.403 ± 0.008 | 0.462 ± 0.009 |
| Surro-seq | CRISMER | **0.718 ± 0.012** | 0.287 ± 0.010 | **0.466 ± 0.018** |
| | CRISPR-DIPOFF [31] | 0.667 ± 0.011 | 0.330 ± 0.014 | 0.310 ± 0.015 |
| | CRISOT [5] | 0.709 ± 0.012 | **0.432 ± 0.017** | 0.458 ± 0.018 |
| | CRISPR-BERT [23] | 0.712 ± 0.011 | 0.367 ± 0.012 | 0.376 ± 0.019 |
| | CCLMoff [13] | 0.705 ± 0.012 | 0.411 ± 0.015 | 0.437 ± 0.018 |
| GUIDE-seq | CRISMER | **0.993 ± 0.002** | 0.080 ± 0.004 | **0.526 ± 0.023** |
| | CRISPR-DIPOFF [31] | 0.953 ± 0.006 | 0.225 ± 0.013 | 0.190 ± 0.015 |
| | CRISOT [5] | 0.990 ± 0.003 | **0.346 ± 0.016** | 0.380 ± 0.022 |
| | CRISPR-BERT [23] | 0.993 ± 0.002 | 0.196 ± 0.009 | 0.358 ± 0.025 |
| | CCLMoff [13] | 0.986 ± 0.003 | 0.192 ± 0.010 | 0.288 ± 0.021 |
| TTISS | CRISMER | 0.977 ± 0.003 | 0.121 ± 0.004 | **0.436 ± 0.016** |
| | CRISPR-DIPOFF [31] | 0.896 ± 0.007 | 0.169 ± 0.009 | 0.122 ± 0.009 |
| | CRISOT [5] | **0.981 ± 0.002** | **0.465 ± 0.015** | 0.403 ± 0.016 |
| | CRISPR-BERT [23] | 0.971 ± 0.003 | 0.284 ± 0.009 | 0.341 ± 0.018 |
| | CCLMoff [13] | 0.971 ± 0.003 | 0.241 ± 0.009 | 0.257 ± 0.014 |

#### 3.1.3. Testing Scenario 3: Mismatch-and-Indel (Bulge) Test

The first two scenarios consider substitution (mismatch) off-targets only. To assess generalization to insertions and deletions, we evaluated CRISMER on the Group IV bulge benchmark (Table 1), in which an off-target may contain a single-base bulge in addition to mismatches; here CRISMER uses the 20 × 24 encoding described in Section 2.2. We compared against CRISPR-BERT and CCLMoff, the two baselines that natively support indels, and excluded CRISOT and CRISPR-DIPOFF because they are designed for substitution-only off-targets. As shown in Table 5, CRISMER achieved the best result on all three metrics (AUROC 0.990, F1 0.690, PR-AUC 0.771), ahead of CRISPR-BERT (PR-AUC 0.757) and CCLMoff (PR-AUC 0.669). These results indicate that the directional pairwise encoding and multi-scale convolutional design extend naturally to the indel setting, requiring no architectural change beyond the wider input.

**Table 5:** Performance comparison in Testing Scenario 3 (mismatch-and-indel benchmark, Group IV). Values are mean *±* standard deviation over 1,000 bootstrap resamples. CRISOT and CRISPR-DIPOFF are omitted as they do not model indels; the best mean per column is in bold.

| Model | AUROC | F1 | PR-AUC |
| --- | --- | --- | --- |
| CRISMER | **0.990 ± 0.001** | **0.690 ± 0.011** | **0.771 ± 0.012** |
| CRISPR-BERT [23] | 0.987 ± 0.002 | 0.626 ± 0.011 | 0.757 ± 0.012 |
| CCLMoff [13] | 0.978 ± 0.002 | 0.553 ± 0.011 | 0.669 ± 0.015 |

### 3.2. Ablation Study

To isolate the contribution of each architectural component, we performed a systematic ablation using the Testing Scenario 1 setup as the baseline and varying one design choice at a time. Table 6 reports the results, and Figure 2(c,d) shows the corresponding ROC and precision-recall curves. Four conclusions emerge.

**Table 6:** Ablation study under the Testing Scenario 1 setup. Each variant changes one design choice relative to the full model. Values are mean *±* standard deviation over 1,000 bootstrap resamples; the best mean F1 and PR-AUC are in bold.

| Variant | Change | AUROC | F1 | PR-AUC |
| --- | --- | --- | --- | --- |
| Full model | — | $0.992 \pm 0.004$ | **$0.728 \pm 0.030$** | **$0.818 \pm 0.030$** |
| A | $20 \times 5$ encoding | $0.994 \pm 0.003$ | $0.677 \pm 0.031$ | $0.726 \pm 0.038$ |
| B | $20 \times 4$ encoding | $0.994 \pm 0.003$ | $0.684 \pm 0.031$ | $0.759 \pm 0.036$ |
| C | no attention (CBAM off) | $0.993 \pm 0.003$ | $0.719 \pm 0.030$ | $0.813 \pm 0.031$ |
| D | no transformer | $0.991 \pm 0.005$ | $0.606 \pm 0.032$ | $0.700 \pm 0.045$ |
| E | 3 CNN branches | $0.993 \pm 0.003$ | $0.580 \pm 0.029$ | $0.754 \pm 0.039$ |
| F | 5 CNN branches | $0.992 \pm 0.004$ | $0.623 \pm 0.031$ | $0.734 \pm 0.038$ |
| G | no CNN | $0.989 \pm 0.004$ | $0.630 \pm 0.035$ | $0.632 \pm 0.046$ |

First, both major components are necessary. Removing the transformer block (variant D) reduced PR-AUC from 0.818 to 0.700, and removing the convolutional block (variant G) reduced it further to 0.632—the largest drop—confirming that multi-scale *k*-mer extraction by the convolutional branches is the single most important component, with the transformer contributing a substantial additional gain by modelling long-range dependencies. Second, the directional 20 × 16 encoding is justified: replacing it with a 4-channel (variant B, PR-AUC 0.759) or 5-channel (variant A, 0.726) one-hot scheme degraded performance, directly supporting the encoding choice over conventional alternatives. Third, the four-branch width is a good operating point, as both three branches (variant E, 0.754) and five branches (variant F, 0.734) were inferior to the four-branch design. Fourth, the CBAM attention modules provided a small but consistent improvement (variant C without attention, 0.813, versus 0.818 for the full model). Together, these experiments confirm that the reported performance arises from the combination of components rather than from any single one or from tuning alone.

### 3.3. Computational Efficiency

Table 7 compares the neural models in terms of the number of trainable parameters and the average training time per epoch, measured under the Testing Scenario 1 setup using a single NVIDIA Tesla P100 GPU in the Kaggle environment. CRISMER is the most lightweight of the neural models, with 1.19M trainable parameters—roughly 2.3× fewer than CRISPR-DIPOFF, 8.5× fewer than CRISPR-BERT, and almost two orders of magnitude fewer than the foundation-model CCLMoff (99.6M)—and the shortest per-epoch training time (31.9 s). CRISOT is not included because it is a gradient-boosted tree model over precomputed RNA-DNA interaction features rather than a neural network trained by gradient descent, so trainable-parameter count and per-epoch time are not directly comparable. These figures show that CRISMER attains its accuracy with a markedly smaller and faster model than the transformer-and foundation-model baselines, which is advantageous for genome-wide screening, where large numbers of candidate sites must be scored.

**Table 7:** Model size and training cost of the neural baselines. Trainable parameters and average per-epoch training time were measured under the Testing Scenario 1 setup using a single NVIDIA Tesla P100 GPU in the Kaggle environment. CRISOT is excluded as it is a gradient-boosted tree model rather than a neural network.

| Model | Trainable params | Time/epoch (s) |
| --- | --- | --- |
| CRISMER | 1,190,314 | 31.9 |
| CRISPR-DIPOFF [31] | 2,778,370 | 35.8 |
| CRISPR-BERT [23] | 10,175,277 | 54.0 |
| CCLMoff [13] | 99,561,089 | 731.4 |

### 3.4. Model Interpretation

To gain insights into the deep features learned by our model, we employed Integrated Gradients, an axiomatic attribution method, to interpret the model’s predictions. This technique calculates the gradient of the model’s output with respect to the input features and integrates it along a path from a baseline input to the actual input, yielding a quantitative measure of each feature’s contribution to the prediction.

We applied this method to the model trained on 80% of Group III dataset of Table 1 and computed attribution scores for all test samples with respect to the positive class. For each test sample, the attribution scores were normalized across all features to allow consistent comparison. We then averaged the normalized attribution scores for each feature across all test samples to obtain a global importance ranking. Features were ranked based on these averaged normalized scores to highlight their relative contributions to the model’s predictions.

The top 15 features driving positive, negative, and overall predictions are shown in Figure 3a–c. Because the dataset is skewed toward negative examples, the negative-and overall-prediction rankings largely coincide; separating them here (Figure 3b vs. 3c) highlights any subtle differences.

- The most important features correspond to mismatch types; this is expected because mismatches are a key factor in off-target effects. In our study, for 23 positions and four matching base pairs, there are 92 possible matching features and 276 possible mismatching features. In the test set, we observed 86 matching features and 258 mismatching features.
- Based on feature rankings derived from attribution scores, we found that all top 50 features contributing to positive predictions were mismatch-type features. Similarly, for both negative and overall predictions, all top 100 ranked features were mismatch-type features.

**Figure 3:**
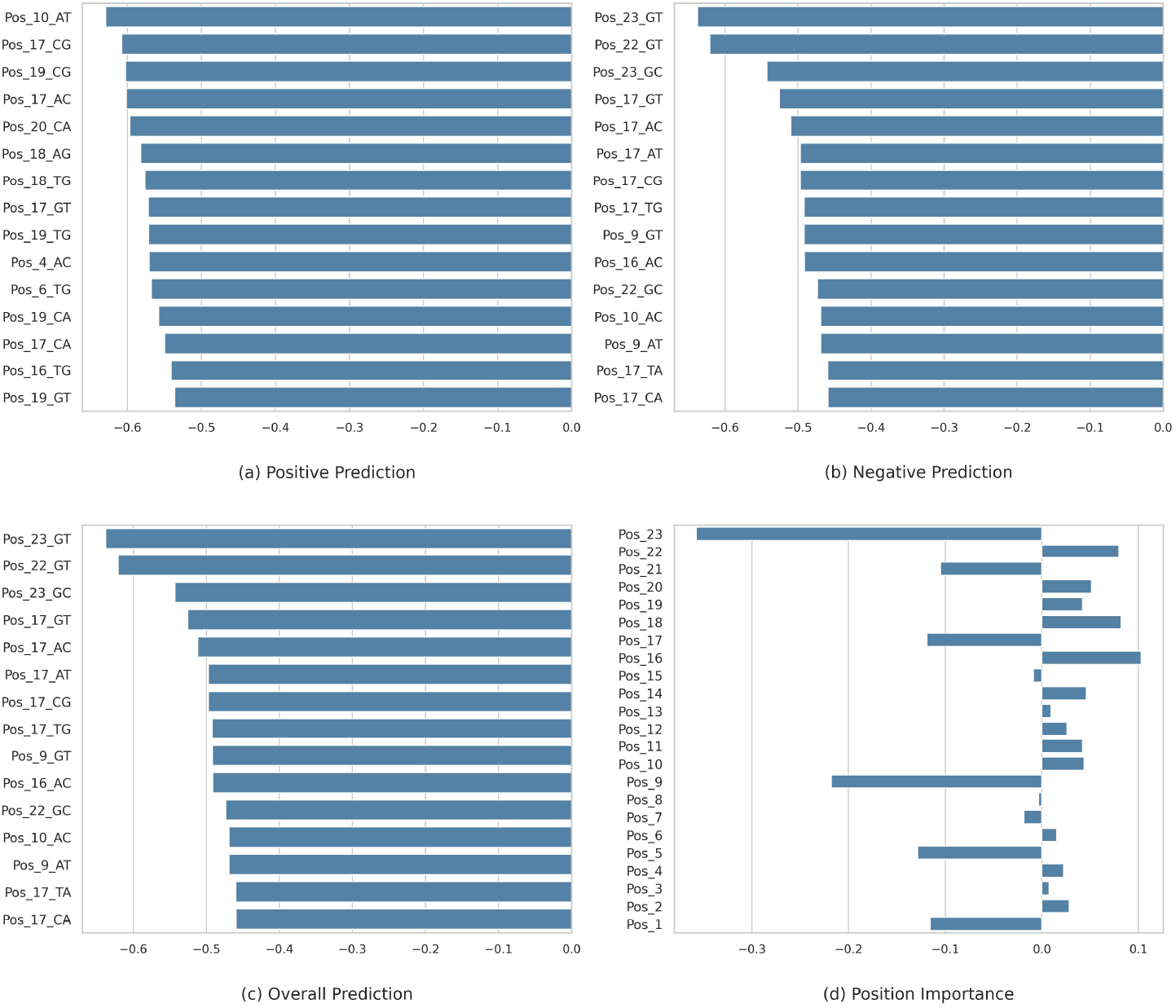
Attribution of feature and position importance for the positive class predictions. (a) Top 15 features contributing to positive predictions, ranked by attribution score. (b) Top 15 features contributing to negative predictions, ranked by attribution score. (c) Top 15 features contributing to overall predictions, ranked by attribution score. (d) Mean importance of each sequence position, providing a position-level perspective of relative contributions to the model’s positive class predictions.

To assess the positional impact of features, we computed the average attribution score for each position. The results are illustrated in Figure 3d, which highlights the key positions influencing model predictions. Our results indicate that PAM-proximal positions contribute more significantly to model predictions than PAM-distal positions. However, some PAM-distal positions, such as positions 1, 5, and 9, also exhibit strong influence, suggesting their potential role in model decision-making.

### 3.5. Efficiency Analysis Results

To evaluate our confidence score, we analyzed the correlation between confidence score and the empirical probability of an off-target being active on the held-out Group II dataset (Table 1; see Methods for calibration and binning details). Figure 4 shows this relationship across score bins for varying softmax temperature parameters (*T* = 1, 5, 10). Active-off-target probability increased with confidence score at all temperature settings, confirming a positive correlation. As expected, lower *T* produced sharper, more concentrated score distributions, while higher *T* produced smoother, more diffuse ones. We set *T* = 10 for the remainder of the study.

**Figure 4:**
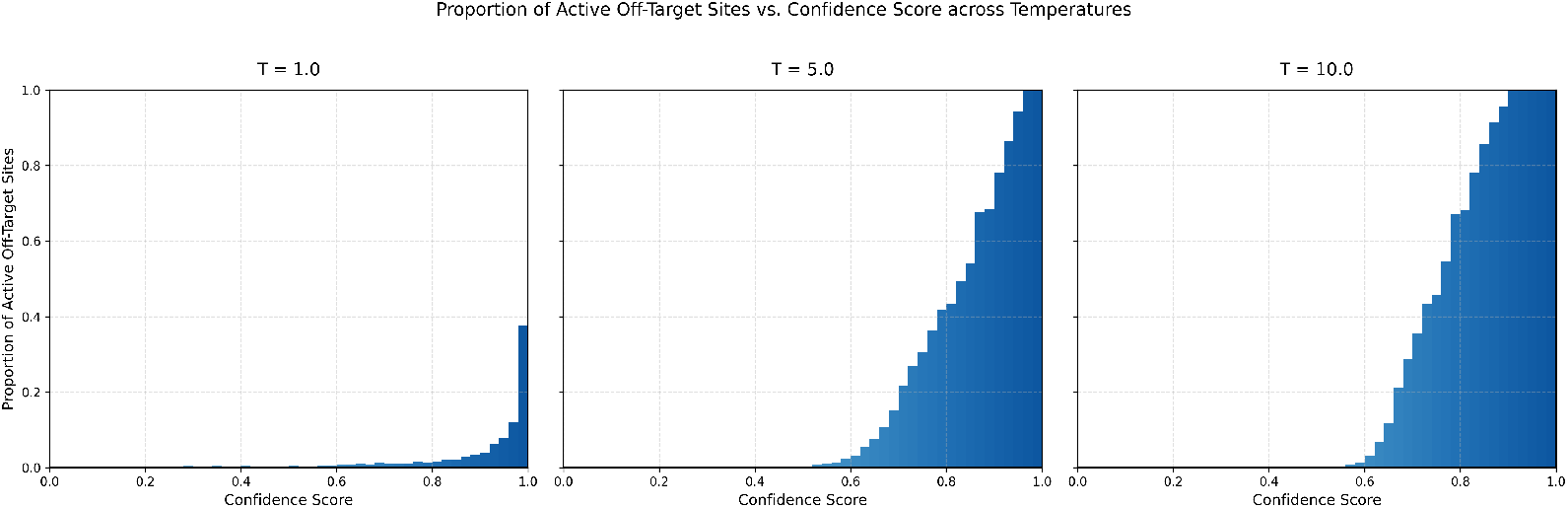
Proportion of active off-target sites across different CRISMER confidence score ranges, based on the model’s final softmax output. Results are averaged over the Group II benchmark datasets.

To confirm this calibration generalizes to unseen data, we repeated an 8-fold rotation on Group II, holding out one fold at a time. The calibration threshold (defined as the score at which empirical active-off-target probability first reaches 80%) was highly stable across folds (0.788 *±* 0.002), and holdout precision at this threshold closely matched the 80% target (0.797 *±* 0.027); Supplementary Table S7, indicating the threshold is not an artifact of a particular data split.

We further compared our model’s calibration behavior against other state-of-the-art off-target prediction methods, applying the identical calibration procedure — including the same data splits — to each method for a fair comparison (Supplementary Figure S4). Our model’s active-off-target probability increased monotonically with confidence score, approaching 1 above a score of 0.9. Other methods showed no comparable monotonic trend and failed to reach similarly high probabilities even in their highest-confidence bins, indicating that, despite competitive discriminative performance, their raw scores are poorly calibrated and not reliably interpretable as activity probabilities.

### 3.6. Comparison of CRISMER-Spec with State-of-the-Art Specificity Evaluation Methods

We evaluated the performance of the CRISMER-Spec specificity score by analyzing its correlation with the number of experimentally validated active off-targets in both the Group-I and Group-II data of Table 1. As shown in Figure 5a, Spearman’s rank correlation coefficient is negative, indicating that higher CRISMER-Spec scores are associated with fewer active off-targets. A stronger (more negative) correlation implies a better ability to distinguish highly specific sgRNAs.

**Figure 5:**
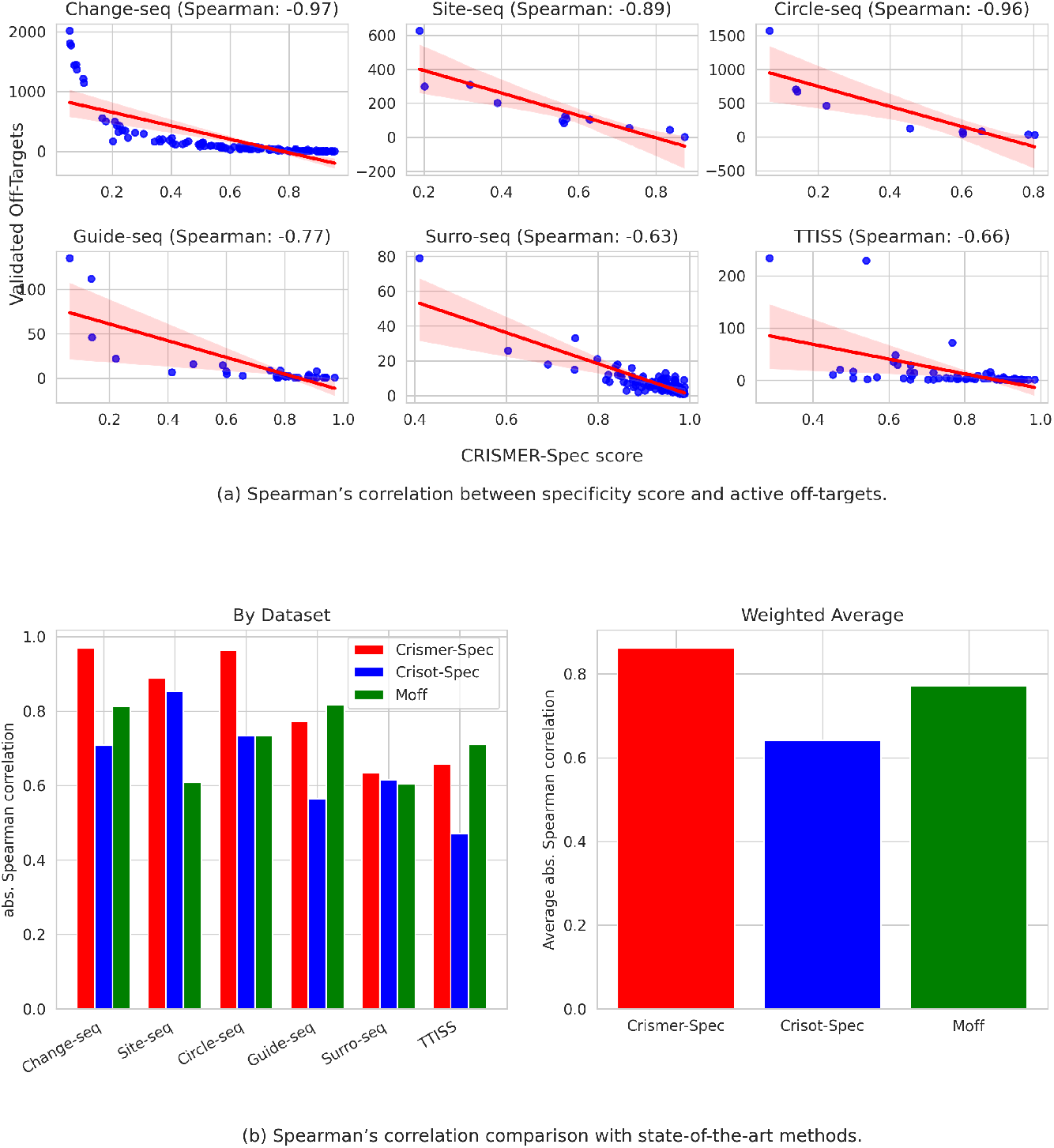
Evaluation of the proposed specificity assessment method. (a) Spearman’s correlation between CRISMER-Spec specificity scores and experimentally validated counts of active off-target sites reported in different studies (Groups I and II benchmark datasets). (b) Comparison of CRISMER-Spec performance against existing sgRNA specificity prediction tools in estimating the number of active off-target sites. Absolute Spearman’s correlation values are reported. The total number of data points in each dataset is used as weights for averaging.

Our method outperforms existing approaches, including CRISOT-Spec and MOFF-aggregate, in terms of absolute Spearman’s correlation between the predicted specificity score and the number of active off-targets. Figure 5b provides a comparative analysis, further demonstrating that CRISMER-Spec achieves stronger correlations and thus more effectively captures genome-wide specificity. The results for MOFF-aggregate and CRISOT-Spec are adopted from [5].

### 3.7. sgRNA Optimization Results Targeting PCSK9, BCL11A, and EXM1 Genes

CRISOT developed an optimization methodology, designated CRISOT-Opti, for sgRNA design. This strategy was applied to two therapeutically significant genes, *PCSK9* and *BCL11A*. The optimized sgRNAs were subsequently validated via whole-genome sequencing (WGS) in HEK293T cells. Furthermore, this approach was extended to the *EXM1* gene and experimentally validated using GUIDE-seq analysis.

In the present study, we optimized sgRNAs for these target genes in a similar manner. Variants exhibiting the highest specificity scores while satisfying the confidence threshold (CRISMER confidence score *≥* 0.85) are shown in Table 8. This threshold was selected because, at this value, more than 80% of off-target sites are active when assessed using a comprehensive dataset combining all Group-II (Table 1) datasets. This study was conducted entirely *in silico*, as wet lab validation was beyond the scope of this work. To provide a robust comparison, we reproduced the results of the established CRISOT-Opti method for the three targets. Similarly, we applied our novel optimization method, CRISMER-Opti, to the same set of variants. The detailed results for both methods are presented in Supplementary Data.

**Table 8:**
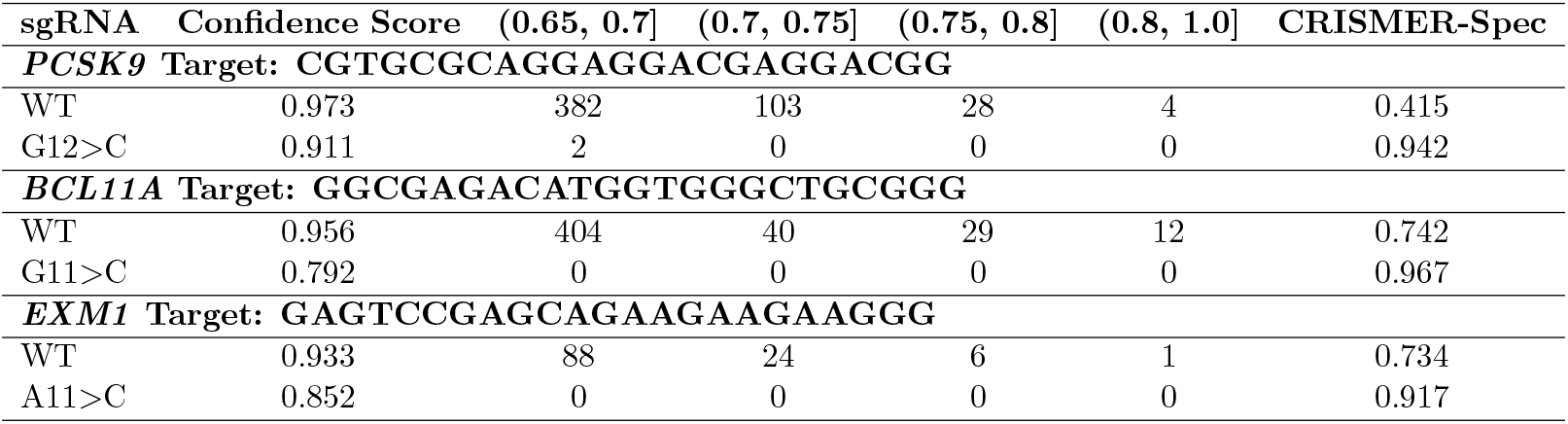
Genome-wide optimization results for *PCSK9*, *BCL11A*, and *EXM1* target sequences. For each wild-type (WT) and optimized sgRNA variant, the table reports the confidence score, the number of predicted off-target sites within specified confidence score intervals, and the CRISMER-Spec specificity score.

Notably, for *BCL11A*, CRISOT reported a G11*>*T substitution that enhanced off-target specificity while maintaining substantial on-target efficiency, as validated by WGS. Here, the notation G11*>*T denotes a point mutation where the guanine (G) at position 11 in the sgRNA sequence is replaced by a thymine (T). Our reproduction of CRISOT-Opti identified three variants, namely, T18*>*C, G11*>*A, and G19*>*A, meeting their optimization threshold (CRISOT-Score *≥* 0.8) with higher specificity scores than G11*>*T; to our knowledge, these three variants have not been experimentally tested, so no conclusion can be drawn about their actual performance. In contrast, our methodology assigned the experimentally validated G11*>*T variant a higher specificity score than these three alternatives. While this single case is consistent with CRISMER-Opti’s ranking better reflecting experimentally relevant off-target behavior, we emphasize that this is an *in silico* observation drawn from one gene target and cannot, on its own, establish general superiority; broader experimental validation across additional targets would be required to support that conclusion. For *EXM1*, our optimized sgRNA was identical to the one experimentally validated using GUIDE-seq in the CRISOT study.

## 4. Conclusion

CRISPR-Cas9 has revolutionized the field of genome editing by enabling precise, efficient, and targeted modifications to genetic material. Despite its transformative potential, off-target effects remain a major limitation, raising concerns about the safety and reliability of CRISPR-based applications, particularly in therapeutic contexts. To address these challenges, computational models have emerged as indispensable tools for improving guide RNA (sgRNA) design, enhancing on-target efficiency, and minimizing off-target activity.

In this study, we presented a hybrid deep learning architecture that integrates multi-branch Convolutional Neural Networks (CNNs) with multi-head attention mechanisms inspired by transformer models. This design effectively captures both local sequence patterns and long-range dependencies, yielding superior performance in CRISPR off-target prediction compared to existing approaches. A systematic ablation study confirmed that each component—the multi-scale convolutional branches, the transformer block, and the directional sparse encoding—contributes to the final performance, and the model generalized to both fully independent datasets and a mismatch-and-indel benchmark while remaining more compact and faster to train than the transformer-and foundation-model baselines. A key feature of our model is its emphasis on interpretability. This not only provides greater transparency in predictive outcomes but also aids researchers in understanding the factors influencing sgRNA activity and specificity.

Beyond off-target prediction, we extended our study to optimize single guide RNA (sgRNA) sequences. We identified optimized sgRNAs targeting the *PCSK9*, *BCL11A*, and *EXM1* genes. For *EXM1*, our result aligns with the experimentally validated sgRNA reported in CRISOT. In the case of *BCL11A*, our method prioritizes the experimentally validated sgRNA over the one reported in CRISOT. Future experimental validation can be carried out for additional sgRNAs identified in this study, as well as for those targeting other genes.

We reiterate that the sgRNA optimization results presented in this work are computational predictions. While they are grounded in a calibrated confidence score and, where possible, benchmarked against previously published experimental outcomes, they have not yet been validated experimentally in this study. We further note that the present optimization experiments considered mismatch-type sequence variants only; indel-based variants were not included in the sgRNA search space. Extending the optimization framework to incorporate indels and pursuing experimental validation of the proposed variants remain important directions for future work.

Overall, our work contributes a robust and interpretable solution to CRISPR off-target prediction and sgRNA optimization. By combining deep learning innovations with practical applications, we take a step closer to safer, more accurate genome editing technologies. Future efforts should focus on expanding model capabilities, incorporating additional biological features, and validating computational predictions through experimental assays to ensure real-world effectiveness.

## 5. Author contributions statement

Rahman perceived the research work. Emtiaj and Rafi contributed equally in developing the methods, experimenting, and analyzing and interpreting the results. Rahman and Nayeem verified the analysis and results. Emtiaj and Rafi wrote the first draft of the manuscript. Rahman and Nayeem supervised the project and contributed to finalizing the manuscript.

## Acknowledgments

The authors thank the anonymous reviewers for their valuable suggestions. This work was supported by the Research and Innovation Centre for Science and Engineering at BUET (RISE-BUET) Internal Research Grant (ID: 2024-01-084).

## Notes

https://github.com/Anamul-Hoque-Emtiaj/CRISMER

### Competing Interest Statement

The authors have declared no competing interest.

### Summary of Updates

Included an ablation study, incorporated a comparison with one additional benchmark study, and performed a new experiment on the indel dataset.

