## Supplementary Data 1 for "CRISMER: A Transformer-based Interpretable Deep Learning Approach for Genome-wide CRISPR-Cas9 Off-Target Prediction and Optimization"

**S3 ROC, Precision-Recall, and Training Loss Convergence Curves** **7**

**S4 Calibration and Threshold Analysis** **8**

**S5 Supplementary Data Files** **9**

S5.1 Supplementary Data 2 (provided as a separate Excel file: SD-2.xlsx) . . . . . . . 9

### S1 Hyperparameter Optimization and Search Space

To optimize the hyperparameters of CRISMER, we utilized a genetic-algorithm (GA) search. The population size was set to 20 and the search was run for 5 generations. The optimization search space of the genetic algorithm is shown in Table S1. The final optimized hyperparameters for Testing Scenario 1, Testing Scenario 2, and Testing Scenario 3 are provided in Table S2.

Table S1: Genetic Algorithm hyperparameter optimization search space.

| **Hyperparameter** | **Description** | **Search Space Values** |
|---|---|---|
| num_layers | Number of Transformer encoder layers in the model architecture. | [1, 2, 3] |
| num_heads | Number of attention heads in the multi-head attention mechanism. | [4, 8, 16] |
| number_hidder_layers | Number of fully connected hidden layers after the Transformer module. | [1, 2] |
| dropout_prob | Dropout probability used in the Transformer layers and feedforward hidden layers. | [0.1, 0.2, 0.3] |
| batch_size | The number of training samples processed in one forward/backward pass. | [32, 64, 128] |
| epochs | Total number of complete passes through the training dataset. | [30, 40, 50, 80] |
| learning_rate | Step size optimization parameter for the Adam optimizer. | [1e-5, 5e-5, 1e-4, 1e-3, 5e-3] |
| pos_weight | Weight coefficient applied to positive class targets to handle class imbalance. | [5, 8, 10, 20, 30, 50] |
| attn | Flag indicating whether Channel and Spatial Attention mechanisms are enabled. | [True, False] |

Table S2: Optimized hyperparameters selected by the genetic algorithm for each testing scenario.

| **Hyperparameter** | **Description** | **Scenario 1** | **Scenario 2** | **Scenario 3** |
|---|---|---|---|---|
| num_layers | Number of Transformer encoder layers in the model architecture. | 2 | 2 | 2 |
| num_heads | Number of attention heads in the multi-head attention mechanism. | 4 | 4 | 8 |
| number_hidder_layers | Number of fully connected hidden layers after the Transformer module. | 1 | 2 | 1 |
| dropout_prob | Dropout probability used in the Transformer layers and feedforward hidden layers. | 0.2 | 0.2 | 0.2 |
| batch_size | The number of training samples processed in one forward/backward pass. | 64 | 128 | 32 |
| epochs | Total number of complete passes through the training dataset. | 40 | 50 | 40 |
| learning_rate | Step size optimization parameter for the Adam optimizer. | 0.0001 | 0.001 | 5e-05 |
| pos_weight | Weight coefficient applied to positive class targets to handle class imbalance. | 10 | 30 | 8 |
| attn | Flag indicating whether Channel and Spatial Attention mechanisms are enabled. | True | False | True |

### S2 Model Performance with 95% Confidence Intervals

In this section, we present the complete tables showing the performance metrics of all models under all scenarios. While the main text reports only the mean ± standard deviation, here we report both the mean ± standard deviation and the corresponding 95% percentile bootstrap confidence intervals (CI) over 1,000 bootstrap resamples. For each metric cell, the top line reports the Mean ± SD and the bottom line reports the 95% bootstrap confidence interval in brackets.

#### S2.1 Testing Scenario 1

Testing Scenario 1 replicates the experimental setup of CRISPR-DIPOFF using the Group III dataset, with an 80%/20% random split. The results are reported in Table S3.

Table S3: Testing Scenario 1 performance summary with 95% bootstrap confidence intervals.

| Model | ROC AUC | F1-Score | PR AUC |
|---|---|---|---|
| **CRISMER** | $0.9923 \pm 0.0039$<br>$[0.9838, 0.9980]$ | $0.7279 \pm 0.0298$<br>$[0.6667, 0.7867]$ | $0.8177 \pm 0.0301$<br>$[0.7530, 0.8728]$ |
| CRISPR-DipOff | $0.9930 \pm 0.0020$<br>$[0.9889, 0.9963]$ | $0.6951 \pm 0.0324$<br>$[0.6271, 0.7539]$ | $0.7175 \pm 0.0382$<br>$[0.6367, 0.7874]$ |
| CRISOT | $0.9931 \pm 0.0031$<br>$[0.9861, 0.9979]$ | $0.6530 \pm 0.0380$<br>$[0.5744, 0.7228]$ | $0.7486 \pm 0.0380$<br>$[0.6692, 0.8179]$ |
| CRISPR-BERT | $0.9930 \pm 0.0029$<br>$[0.9864, 0.9972]$ | $0.6218 \pm 0.0389$<br>$[0.5463, 0.6965]$ | $0.7174 \pm 0.0392$<br>$[0.6423, 0.7892]$ |
| CCLMoff | $0.9782 \pm 0.0037$<br>$[0.9702, 0.9852]$ | $0.3427 \pm 0.0285$<br>$[0.2869, 0.3974]$ | $0.4588 \pm 0.0439$<br>$[0.3734, 0.5482]$ |

#### S2.2 Testing Scenario 2

Testing Scenario 2 represents the independent dataset evaluation. Models were trained on Group I (CHANGE-seq + SITE-seq) and evaluated on Group II datasets (CIRCLE-seq, Surro-seq, GUIDE-seq, and TTISS). The results are reported in Table S4.

Table S4: Testing Scenario 2 performance summary with 95% bootstrap confidence intervals.

| Dataset | Model | ROC AUC | F1-Score | PR AUC |
|---|---|---|---|---|
| **Circleseq** | **CRISMER** | $0.9725 \pm 0.0012$ [0.9700, 0.9750] | $0.5603 \pm 0.0057$ [0.5491, 0.5715] | $0.6541 \pm 0.0080$ [0.6386, 0.6693] |
| | CRISPR-DipOff | $0.8686 \pm 0.0032$ [0.8623, 0.8749] | $0.2605 \pm 0.0082$ [0.2451, 0.2766] | $0.3459 \pm 0.0085$ [0.3292, 0.3613] |
| | CRISOT | $0.9529 \pm 0.0015$ [0.9498, 0.9558] | $0.2528 \pm 0.0085$ [0.2352, 0.2679] | $0.4985 \pm 0.0087$ [0.4811, 0.5154] |
| | CRISPR-BERT | $0.9715 \pm 0.0011$ [0.9693, 0.9735] | $0.5729 \pm 0.0071$ [0.5593, 0.5870] | $0.6201 \pm 0.0084$ [0.6035, 0.6359] |
| | CCLMoff | $0.9485 \pm 0.0016$ [0.9453, 0.9515] | $0.4025 \pm 0.0084$ [0.3859, 0.4195] | $0.4622 \pm 0.0088$ [0.4448, 0.4793] |
| **surroseq** | **CRISMER** | $0.7179 \pm 0.0115$ [0.6955, 0.7396] | $0.2872 \pm 0.0095$ [0.2682, 0.3055] | $0.4656 \pm 0.0175$ [0.4309, 0.5002] |
| | CRISPR-DipOff | $0.6671 \pm 0.0112$ [0.6443, 0.6884] | $0.3295 \pm 0.0135$ [0.3035, 0.3561] | $0.3103 \pm 0.0146$ [0.2829, 0.3403] |
| | CRISOT | $0.7092 \pm 0.0116$ [0.6861, 0.7315] | $0.4321 \pm 0.0172$ [0.3988, 0.4651] | $0.4577 \pm 0.0175$ [0.4241, 0.4916] |
| | CRISPR-BERT | $0.7124 \pm 0.0112$ [0.6904, 0.7344] | $0.3671 \pm 0.0124$ [0.3443, 0.3922] | $0.3755 \pm 0.0189$ [0.3410, 0.4150] |
| | CCLMoff | $0.7048 \pm 0.0115$ [0.6829, 0.7257] | $0.4107 \pm 0.0146$ [0.3816, 0.4378] | $0.4367 \pm 0.0175$ [0.4016, 0.4688] |
| **guideseq** | **CRISMER** | $0.9929 \pm 0.0023$ [0.9875, 0.9965] | $0.0796 \pm 0.0037$ [0.0723, 0.0867] | $0.5261 \pm 0.0229$ [0.4797, 0.5697] |
| | CRISPR-DipOff | $0.9530 \pm 0.0057$ [0.9412, 0.9638] | $0.2246 \pm 0.0133$ [0.1975, 0.2509] | $0.1902 \pm 0.0154$ [0.1609, 0.2213] |
| | CRISOT | $0.9897 \pm 0.0029$ [0.9835, 0.9946] | $0.3460 \pm 0.0161$ [0.3151, 0.3778] | $0.3802 \pm 0.0219$ [0.3363, 0.4215] |
| | CRISPR-BERT | $0.9929 \pm 0.0021$ [0.9879, 0.9963] | $0.1955 \pm 0.0085$ [0.1792, 0.2119] | $0.3583 \pm 0.0248$ [0.3105, 0.4058] |
| | CCLMoff | $0.9860 \pm 0.0025$ [0.9804, 0.9904] | $0.1922 \pm 0.0095$ [0.1733, 0.2106] | $0.2883 \pm 0.0210$ [0.2486, 0.3290] |
| **ttiss** | **CRISMER** | $0.9769 \pm 0.0027$ [0.9715, 0.9817] | $0.1207 \pm 0.0040$ [0.1133, 0.1290] | $0.4361 \pm 0.0158$ [0.4051, 0.4685] |
| | CRISPR-DipOff | $0.8964 \pm 0.0069$ [0.8824, 0.9096] | $0.1690 \pm 0.0094$ [0.1506, 0.1887] | $0.1221 \pm 0.0092$ [0.1051, 0.1413] |
| | CRISOT | $0.9814 \pm 0.0020$ [0.9770, 0.9851] | $0.4652 \pm 0.0151$ [0.4373, 0.4948] | $0.4028 \pm 0.0159$ [0.3723, 0.4351] |
| | CRISPR-BERT | $0.9714 \pm 0.0025$ [0.9664, 0.9761] | $0.2840 \pm 0.0093$ [0.2663, 0.3022] | $0.3412 \pm 0.0182$ [0.3040, 0.3771] |
| | CCLMoff | $0.9714 \pm 0.0029$ [0.9653, 0.9767] | $0.2410 \pm 0.0088$ [0.2235, 0.2593] | $0.2569 \pm 0.0142$ [0.2303, 0.2858] |

#### S2.3 Testing Scenario 3

Testing Scenario 3 evaluates generalization to off-targets containing single-base indels (bulges) using the Group IV benchmark. The results are reported in Table S5.

Table S5: Testing Scenario 3 performance summary with 95% bootstrap confidence intervals.

| Model | ROC AUC | F1-Score | PR AUC |
|---|---|---|---|
| **CRISMER** | $0.9897 \pm 0.0010$ $[0.9878, 0.9915]$ | $0.6904 \pm 0.0109$ $[0.6687, 0.7113]$ | $0.7705 \pm 0.0119$ $[0.7458, 0.7919]$ |
| CRISPR-BERT | $0.9871 \pm 0.0015$ $[0.9842, 0.9899]$ | $0.6262 \pm 0.0108$ $[0.6056, 0.6473]$ | $0.7573 \pm 0.0122$ $[0.7333, 0.7809]$ |
| CCLMoff | $0.9783 \pm 0.0017$ $[0.9749, 0.9815]$ | $0.5533 \pm 0.0108$ $[0.5323, 0.5738]$ | $0.6692 \pm 0.0146$ $[0.6412, 0.6972]$ |

#### S2.4 Ablation Study

The performance of the various model variants under the component-wise ablation study is reported in Table S6.

Table S6: Ablation study performance summary with 95% bootstrap confidence intervals.

| Model / Variant | Description | ROC AUC | F1-Score | PR AUC |
|---|---|---|---|---|
| **CRISMER (Full Model)** | Full baseline architecture (TS1 baseline) | $0.9923 \pm 0.0039$ $[0.9838, 0.9980]$ | $0.7279 \pm 0.0298$ $[0.6667, 0.7867]$ | $0.8177 \pm 0.0301$ $[0.7530, 0.8728]$ |
| Ablation A | 20x5 encoder configuration (in_channels=5) | $0.9940 \pm 0.0025$ $[0.9881, 0.9976]$ | $0.6772 \pm 0.0306$ $[0.6143, 0.7362]$ | $0.7264 \pm 0.0375$ $[0.6506, 0.7955]$ |
| Ablation B | 20x4 encoder configuration (in_channels=4) | $0.9937 \pm 0.0027$ $[0.9876, 0.9978]$ | $0.6844 \pm 0.0310$ $[0.6202, 0.7405]$ | $0.7593 \pm 0.0358$ $[0.6853, 0.8255]$ |
| Ablation C | Channel/Spatial Attention disabled (attn=False) | $0.9933 \pm 0.0033$ $[0.9857, 0.9984]$ | $0.7191 \pm 0.0295$ $[0.6612, 0.7754]$ | $0.8132 \pm 0.0306$ $[0.7491, 0.8676]$ |
| Ablation D | Exclude Transformer module | $0.9908 \pm 0.0047$ $[0.9798, 0.9974]$ | $0.6062 \pm 0.0315$ $[0.5427, 0.6667]$ | $0.6999 \pm 0.0447$ $[0.6130, 0.7867]$ |
| Ablation E | CNN with 3 branches (instead of 4) | $0.9931 \pm 0.0028$ $[0.9869, 0.9974]$ | $0.5804 \pm 0.0289$ $[0.5223, 0.6359]$ | $0.7543 \pm 0.0388$ $[0.6753, 0.8247]$ |
| Ablation F | CNN with 5 branches (instead of 4) | $0.9915 \pm 0.0035$ $[0.9836, 0.9972]$ | $0.6229 \pm 0.0313$ $[0.5607, 0.6798]$ | $0.7339 \pm 0.0379$ $[0.6559, 0.8019]$ |
| Ablation G | Exclude CNN module | $0.9890 \pm 0.0037$ $[0.9805, 0.9950]$ | $0.6304 \pm 0.0351$ $[0.5568, 0.6962]$ | $0.6321 \pm 0.0464$ $[0.5362, 0.7195]$ |

#### S3 ROC, Precision-Recall, and Training Loss Convergence Curves

This section contains the ROC curves, Precision-Recall curves, and training loss convergence curves generated from our evaluation. Each figure presents three panels: (A) the Receiver Operating Characteristic (ROC) curve with per-model AUROC, (B) the Precision-Recall curve with per-model AUPRC, and (C) the cross-entropy training loss versus epoch, illustrating convergence behavior. Figure S1 shows these curves for Testing Scenario 1. Figure S2 shows them for Testing Scenario 3. Figure S3 shows them for the ablation study.

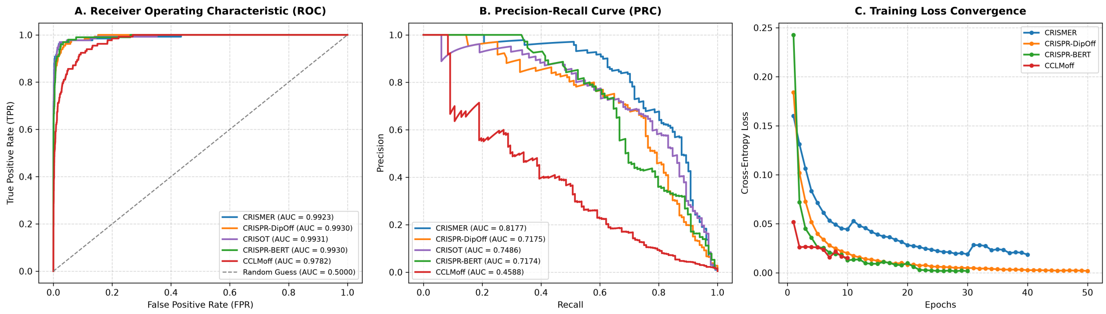

Figure S1: Testing Scenario 1: (A) ROC curves, (B) Precision-Recall curves, and (C) training loss convergence across epochs for all evaluated models.

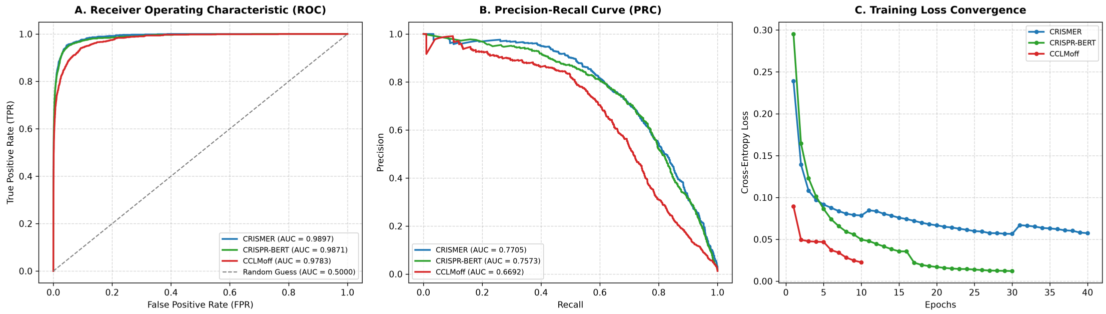

Figure S2: Testing Scenario 3 (mismatch-and-indel bulge setting): (A) ROC curves, (B) Precision-Recall curves, and (C) training loss convergence across epochs for all evaluated models.

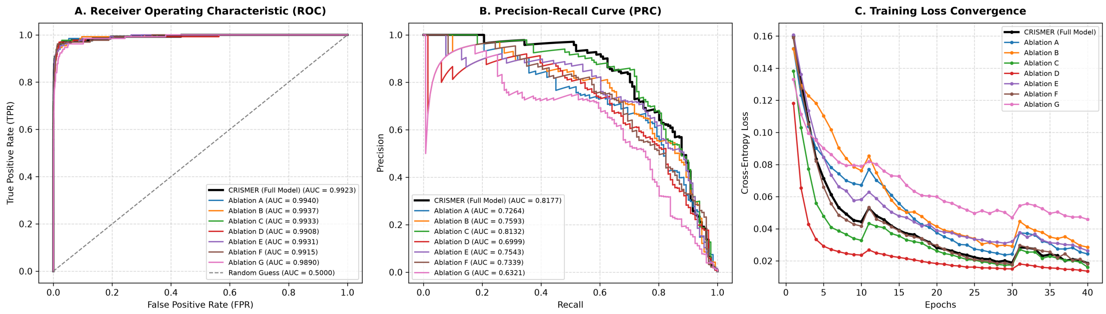

Figure S3: Ablation Study: (A) ROC curves, (B) Precision-Recall curves, and (C) training loss convergence across epochs for the full model and all ablation variants.

### S4 Calibration and Threshold Analysis

To establish a calibrated threshold for active off-target classification and to avoid circularity (leakage) between threshold derivation and evaluation, we performed an 8-fold cross-validation rotation on the Group II dataset. The Group II dataset is entirely held out from model training. The threshold was defined as the confidence score at which the empirical active-off-target probability first reaches 80% on the calibration fold, and evaluated on the holdout fold. Table S7 reports the results of this rotation.

Figure S4 shows the calibration curves for the baseline prediction methods: CRISPR-DIPOFF, CRISPR-BERT, and CCLMoff. These models show a clear lack of calibration and monotonic scaling with empirical off-target cleavage probability compared to CRISMER (whose calibrated monotonic response is detailed in the main manuscript).

Table S7: 8-fold cross-validation threshold calibration and validation on Group II.

| Fold | Threshold | Calib Active Ratio | Holdout Active Ratio | N Holdout Sites | Holdout Precision |
|---|---|---|---|---|---|
| 0 | 0.786208 | 0.006165 | 0.006164 | 114058 | 0.7746 |
| 1 | 0.785840 | 0.006165 | 0.006164 | 114058 | 0.7518 |
| 2 | 0.786286 | 0.006165 | 0.006164 | 114058 | 0.7826 |
| 3 | 0.790061 | 0.006165 | 0.006164 | 114058 | 0.8258 |
| 4 | 0.787125 | 0.006165 | 0.006164 | 114058 | 0.7836 |
| 5 | 0.790418 | 0.006165 | 0.006164 | 114058 | 0.8321 |
| 6 | 0.790061 | 0.006164 | 0.006172 | 114058 | 0.8240 |
| 7 | 0.788861 | 0.006165 | 0.006164 | 114057 | 0.8000 |
| **Mean** | **0.788107** | **0.006165** | **0.006165** | **114057** | **0.7968** |
| **Std Dev** | **0.001822** | **0.000000** | **0.000003** | **0** | **0.0268** |
| **Max - Min** | **0.004578** | **0.000001** | **0.000008** | **1** | **0.0803** |

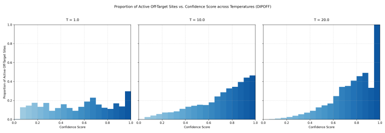

(a) CRISPR-DIPOFF

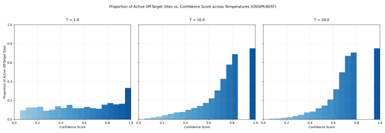

(b) CRISPR-BERT

Figure S4: Cleavage efficiency calibration curves for baseline prediction methods. The empirical proportion of active off-target sites (y-axis) is plotted against model confidence scores (x-axis) grouped into bins.

### S5 Supplementary Data Files

In addition to the tables and figures compiled in this document, the supplementary materials include the following high-throughput optimization dataset:

#### S5.1 Supplementary Data 2 (provided as a separate Excel file: SD-2.xlsx)

This spreadsheet contains the detailed, nucleotide-by-nucleotide in silico optimization results generated by CRISMER-Opti and CRISOT-Opti for three therapeutic target genes:

- **BCL11A:** A gene target relevant for sickle cell disease and $\beta$-thalassemia.
- **EXM1:** A target widely used in human genome engineering studies.
- **PCSK9:** A gene target relevant for hypercholesterolemia.

For each gene target, the spreadsheet lists the sgRNA variant sequence, mutation details (relative to the wild-type), the raw model prediction score, and the aggregated genome-wide specificity score. The sheets in the Excel file are structured as follows:

- **CRISMER-Opti_bcl11a**, **CRISMER-Opti_exm1**, and **CRISMER-Opti_pcsk9**: sgRNA variants designed using the CRISMER model, sorted by their genome-wide specificity score.
- **CRISOT-Opti_bcl11a**, **CRISOT-Opti_exm1**, and **CRISOT-Opti_pcsk9**: Reproductions of the sgRNA variants designed using the CRISOT method.
